# Dorsomedial hypothalamic BDNF neurons integrate thermal afferent signals to control energy expenditure

**DOI:** 10.1101/2021.04.11.439314

**Authors:** Qian Zhou, Hao Bian, Mengting Wang, Xinyan Ni, Wen Z. Yang, Hongbin Sun, Wei L. Shen

## Abstract

Mutations in the gene brain-derived neurotrophic factor (BDNF) cause obesity in humans. BDNF signaling and its expressing neurons in the hypothalamus help control feeding, energy expenditure (EE), and physical activity. However, whether the BDNF neurons interact with another EE-regulating system, the thermoregulation circuitry, remains unclear. Here, we show that BDNF neurons in the dorsomedial hypothalamus (DMH) are activated by environmental cooling and sufficient to induce body temperature increases and brown adipose tissue (BAT) thermogenesis. Conversely, blocking these neurons impairs BAT thermogenesis and cold defense, causing body weight gain and glucose intolerance. DMH BDNF neurons are therefore an important type of thermoregulatory neuron, integrating thermal afferent signals to control EE during cold defense. This reveals a critical intersection between the BDNF circuitry and the thermoregulatory system.

## Introduction

Genes encoding brain-derived neurotrophic factor (*BDNF*) and its receptor, tyrosine receptor kinase B (*TrkB*), have been linked to human obesity, as loss-of-function mutations in either gene cause severe obesity (***Wen et al. 2012; Gray et al. 2006; Yeo et al. 2004***). Studies in rodents have shown that BDNF and its neural circuitry in the hypothalamus control feeding, adaptive thermogenesis, and physical activity (***Xu & Xie 2016; Wang et al. 2020; An et al. 2015; Unger et al. 2007***). For example, genetic deletion of BDNF in the mouse ventromedial hypothalamus (VMH) and the dorsomedial hypothalamus (DMH) causes hyperphagic behavior and obesity (***Unger et al. 2007***). BDNF neurons within the anterior and posterior part of the PVN control feeding and thermogenesis, respectively (***An et al. 2015***). These data suggest that the BDNF circuitry is a crucial player in energy homeostasis. However, whether and how the BDNF circuitry integrates thermal afferent signals to rapidly and efficiently regulate energy expenditure (EE) is unknown. This is a critical gap in our understanding of how BDNF regulates obesity.

Thermal afferent signals are vital for the regulation of body temperature and EE. These signals are delivered via a feed-forward pathway (***Tan & Knight 2018; Morrison & Nakamura 2019***) that begins with thermosensory neurons in the dorsal root ganglion. Information is then relayed by neurons in the spinal cord to the lateral parabrachial nucleus (LPB) where signals are categorized into BAT thermogenesis control signals and vasodilation control signals (***Yang et al. 2020***). Signals are integrated in the preoptic area (POA) and then POA neurons directly target neurons in the DMH or the raphe pallidus nucleus (RPa) to drive different aspects of thermal effector activities, such as brown adipose tissue (BAT) thermogenesis and vasodilation (***Tan & Knight 2018; Morrison & Nakamura 2019***). Blocking Pdyn neurons in the LPB in the warm afferent pathway increases BAT thermogenesis and EE and is protective against weight gain driven by a high-fat diet (***Yang et al. 2020***). Several POA neuron types can induce severe hypothermia and substantially reduce EE (***Morrison & Nakamura 2019; Zhao et al. 2017; Yu et al. 2016***). For example, activation of preoptic neurons that express the leptin receptor reduces core temperature (T_core_) to 30°C and lowers EE by 75% (***Yu et al. 2016***). This also lowers food intake to reduce body weight (***Yu et al. 2016***), which may be explained by lower energy demand in the warm environment. However, whether and how the BDNF circuitry interacts with the thermoregulatory pathway has not been examined.

Here, we found that BDNF neurons in the DMH (DMH^BDNF^) were quickly activated by ambient cold temperatures. Activation of these neurons was sufficient to increase body temperature, EE, and physical activity. Chronic blocking of these neurons impaired glucose metabolism and BAT thermogenesis, leading to weight gain on normal chow. Therefore, our data indicate that DMH^BDNF^ neurons function as a rapid cooling sensor in the thermoregulatory pathway, quickly detecting ambient thermal inputs and then adjusting EE to meet the energy demand required to maintain core temperature (T_core_).

## Results and discussion

### DMH^BDNF^ neurons are sensitive to cold temperature

We sought to investigate whether BDNF neurons in the DMH are sensitive to ambient temperature, as the DMH is considered a center for controlling adaptive thermogenesis (***Tan & Knight 2018; Morrison & Nakamura 2019***). To label BDNF neurons, we crossed BDNF-IRES-Cre mice (***Luo et al. 2019***) to a Cre-dependent GFP reporter line (GFPL10). ∼35% glutamate neurons and ∼15% GABA neurons in the DMH were BDNF-positive, and most of BDNF neurons were glutamatergic (∼60%) (***Figure 1A-C***). BDNF neurons overlapped with those that expressed cFos following cold or warm exposure (***Figure 1D***). Quantification of this overlap revealed that in response to cold, ∼ 34.1% of cFos-positive neurons were BDNF neurons, whereas this overlap was only ∼ 13.3% following warm exposure (***Figure 1E-F***). Expressed in a different way, 22.2% of the BDNF neurons in the DMH responded to cold, whereas only 4.7% responded to warmth (***Figure 1E-F***). These results indicate that DMH^BDNF^ neurons responded preferentially to cold rather than warm temperature.

**Figure 1.**
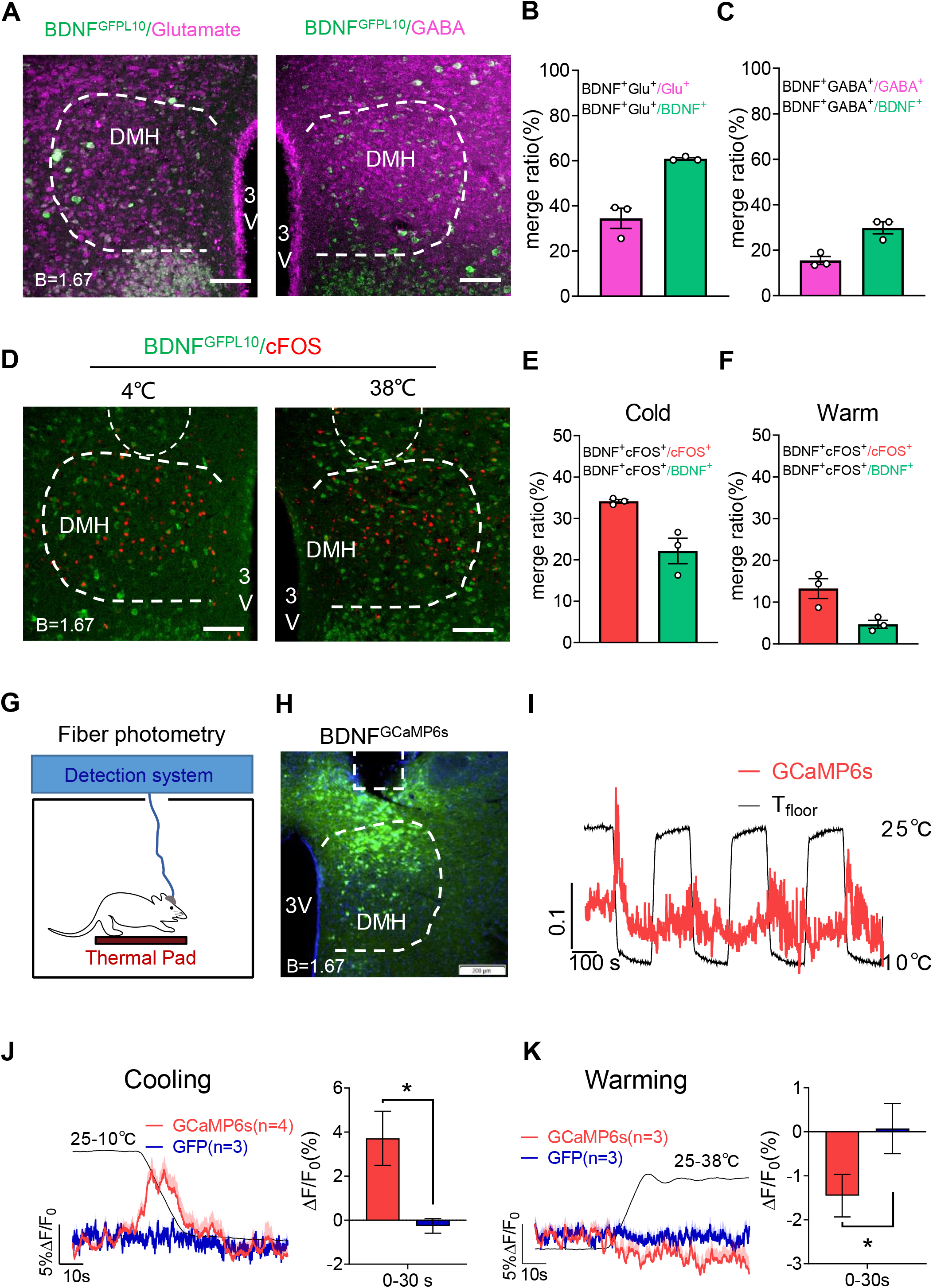
DMH^BDNF^ neurons are sensitive to cold temperature. (**A**) The overlapping between BDNF-IRES-Cre & GFPL10 and glutamate (left panel) or GABA (right panel). Scale bar,100 μm. (**B**) The overlapping rate between BDNF-IRES-Cre & GFPL10 and glutamate in DMH (no. of BDNF^+^ and Glutamate^+^/no. of Glutamate^+^ = 34.5 ± 4.5%, no. of BDNF^+^ and Glutamate^+^/no. of BDNF^+^ = 60.8 ± 0.5%, n = 3). (**C**) The overlapping rate between BDNF-IRES-Cre & GFPL10 and GABA in DMH (no. of BDNF^+^ and GABA^+^/no. of GABA^+^ = 15.5 ± 1.8%, no. of BDNF^+^ and GABA^+^/no. of BDNF^+^ = 29.8 ± 2.7%, n = 3). (**D**) Examples of DMH sections from BDNF-IRES-Cre & GFPL10 mice stained for cFos after cold (4 °C) or warm (38 °C) exposures. Scale bar,100 μm. (**E**) The overlapping between BDNF-IRES-Cre & GFPL10 and cFos induced by cold exposure (4 °C) in DMH (no. of BDNF^+^ and cFos^+^/no. of cFos^+^ = 34.1 ± 0.5%, no. of BDNF^+^ and cFos ^+^/no. of BDNF^+^ = 22.2 ± 3.1%, n = 3). (**F**) The overlapping between BDNF-IRES-Cre & GFPL10 and cFos induced by warm exposure (38 °C) in DMH (no. of BDNF^+^ and cFos^+^/no. of cFos^+^ = 13.3 ±2.4%, no. of BDNF^+^ and cFos^+^/no. of BDNF^+^ = 4.7 ± 1.0%, n = 3). (**G**) Schematic of Ca^2+^ fiber photometry setup. (**H**) Coronal section from BDNF-IRES-Cre mice showing optical fiber path and DIO-GCaMP6s expression. Scale bar, 200 μm. (**I**) Representative GCaMP6s fluorescence trace during cooling cycles. Floor temperature signal trace represents T_floor_ changing from 25 °C to 10 °C. (**J**-**K**) Calcium dynamics from DMH BDNF neurons during cooling (**J**) or warming (**K**) of the floor. ΔF/F_0_ represents the change in GCaMP6s fluorescence from the mean level [t = (−30 to 0 s)]. The GFP was used as a control. Quantification of fluorescence changes post thermal stimuli (0∼30 s) were shown on right panel. B, bregma; 3V, 3rd ventricle; DMH, dorsomedial hypothalamus. All data are shown as means ± SEM. The *P* values were calculated based on two-tailed unpaired t-test, *\*P* ≤ 0.05.

### Calcium dynamics of DMH^BDNF^ neurons to temperature stimuli

To study the speed and dynamics of neural responses to temperature, we injected Cre-dependent AAVs expressing GCaMP6s into the DMH of BDNF-Cre mice (***Figure 1H***), and then used fiber photometry to record calcium activity of DMH^BDNF^ neurons (***Figure 1G-H***) following temperature manipulations. Floor cooling induced a sharp increase in calcium activity in DMH^BDNF^ neurons(***Figure 1I-J***). Interestingly, these responses mainly occurred during the cooling phase, which has also been reported for sensory neurons and the spinal cord (***Yarmolinsky et al. 2016; Wang et al. 2018; Ran et al. 2016***). Once the temperature stopped changing, responses diminished to baseline. This is different from the overall activity of glutamatergic and GABAergic neurons, where there is a plateau of neural activity after temperature stopped changing (***Zhao et al. 2017***). In contrast, warming slightly inhibited the activity of DMH^BDNF^ neurons (***Figure 1K***). These data suggest that DMH^BDNF^ neurons function as a rapid and sensitive detector for cooling, but not for static cold temperatures.

### Activation of DMH^BDNF^ neurons elevates core body temperature, energy expenditure, and physical activity while reducing RER

The selective response to cooling predicts that these BDNF neurons may be necessary for cold-induced thermoregulation. To test this hypothesis, we used the DREADD-hM3Dq system to activate DMH^BDNF^ neurons (***Figure 2A***). As expected, intraperitoneal injection of the hM3Dq agonist CNO (2.5 mg/kg) dramatically increased T_core_, EE, and physical activity (***Figure 2B-D***). The respiratory exchange ratio (RER) dropped significantly following CNO injection (***Figure 2E***), indicating that activation of DMH^BDNF^ neurons increased fat fuel utilization. Daily injection of CNO did not alter food intake (***Figure 2F***), suggesting that DMH^BDNF^ neurons do not regulate feeding. These data indicate that DMH^BDNF^ neurons are sufficient to induce hyperthermia, and elevate EE and physical activity, all of which are consistent with these neurons’ response to cold temperature.

**Figure 2.**
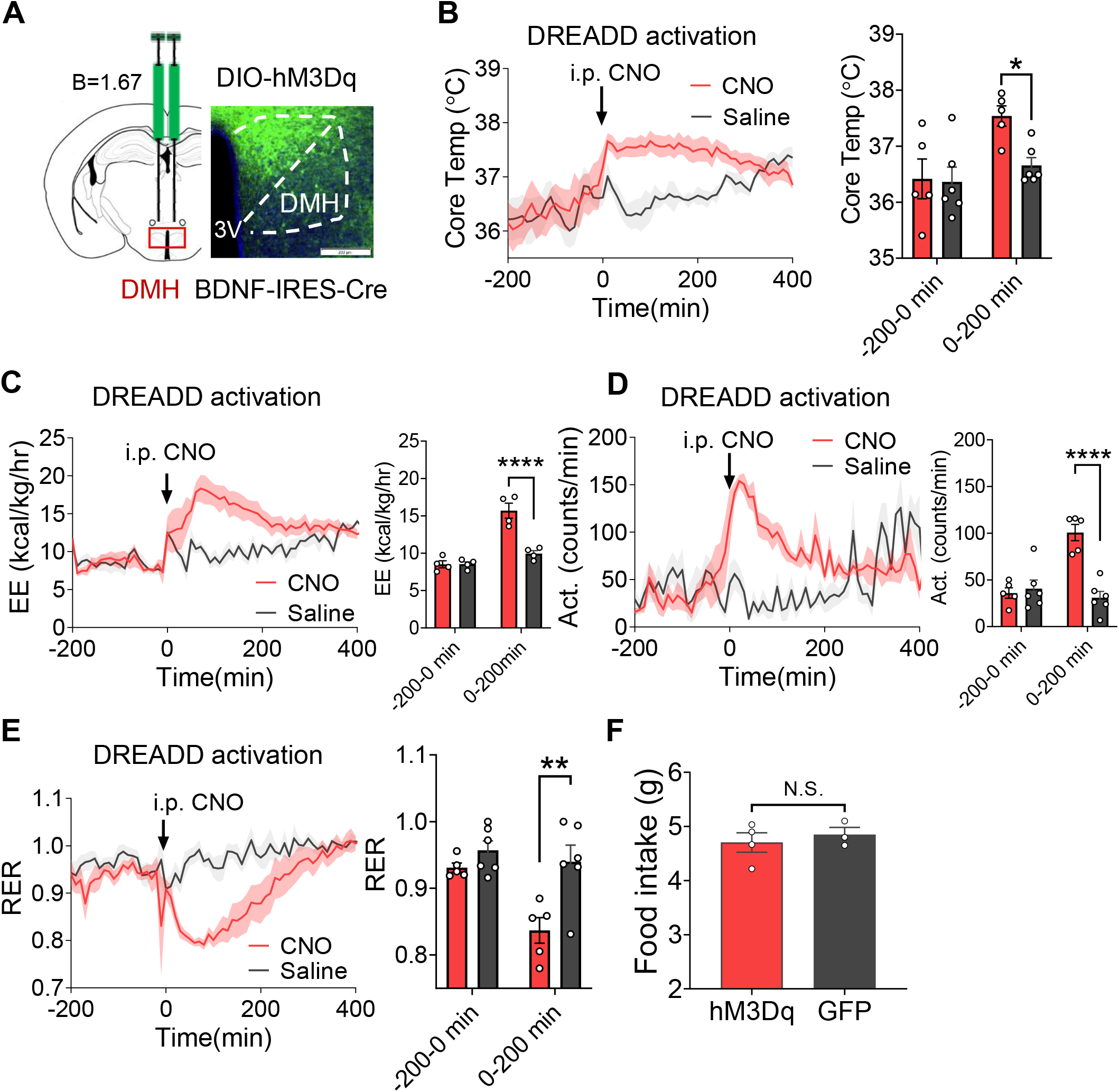
Activation of DMH^BDNF^ neurons elevates core body temperature, energy expenditure, and physical activity while reducing RER. (**A**) Schematic of virus injection (left) and representative expression of hM3Dq-GFP (right) in the DMH of BDNF-IRES-Cre mice, Scale bar, 200 μm. (**B-E**) Core body temperature (**B**), energy expenditure (EE) (**C**), Physical activity (**D**) and Respiratory exchange ratio (RER)(**E**) changes to CNO (i.p, 2.5 mg/kg) or saline in DMH^BDNF & hM3Dq^ mice. For quantification of core body temp, EE, physical activity and RER changes pre (−200-0 min) and post (0-200 min) CNO treatment in hM3Dq and control mice, the bar graph was shown on the right panel. (**F**) Average daily food intake was measured during daily CNO treatment (2.5 mg/kg per day). The *P* values was calculated based on two-tailed unpaired t-test. B, bregma; 3V, 3rd ventricle; DMH, dorsomedial hypothalamus. Except (**F**), the *P* values were calculated on the basis of repeated measures two-way ANOVA with Bonferroni’s corrections. Data are presented as mean± SEM; N.S., not significant, *\*P* ≤ 0.05, *\*\*P* ≤ 0.01, *\*\*\*P* ≤ 0.001 and *\*\*\*\*P* ≤ 0.0001.

### DMH^BDNF^ neurons are required for BAT thermogenesis during cold defense

To test whether DMH^BDNF^ neurons are necessary for thermoregulation and the regulation of EE, we used neural toxin TeNT to block their synaptic transmission (***Figure 3A***). Consistent with DREADD activation studies, mice in which these neurons were chronically inhibited could not maintain T_core_ in a cold environment than GFP controls (***Figure 3B***). Chronic inhibition of DMH^BDNF^ neurons also reduced EE, RER, and physical activity, both in baseline and cold temperatures (***Figure 3C-E***), suggesting long-term effects on both basal metabolism and cold-induced thermogenesis. In contrast, T_core_, EE, RER, and physical activity were not significantly affected in these mice following exposure to warmth (compared to GFP controls) (***Figure 3F-I***). This suggests that blocking DMH^BDNF^ neurons selectively affected thermoregulation, EE, and physical activity in cold defense.

**Figure 3.**
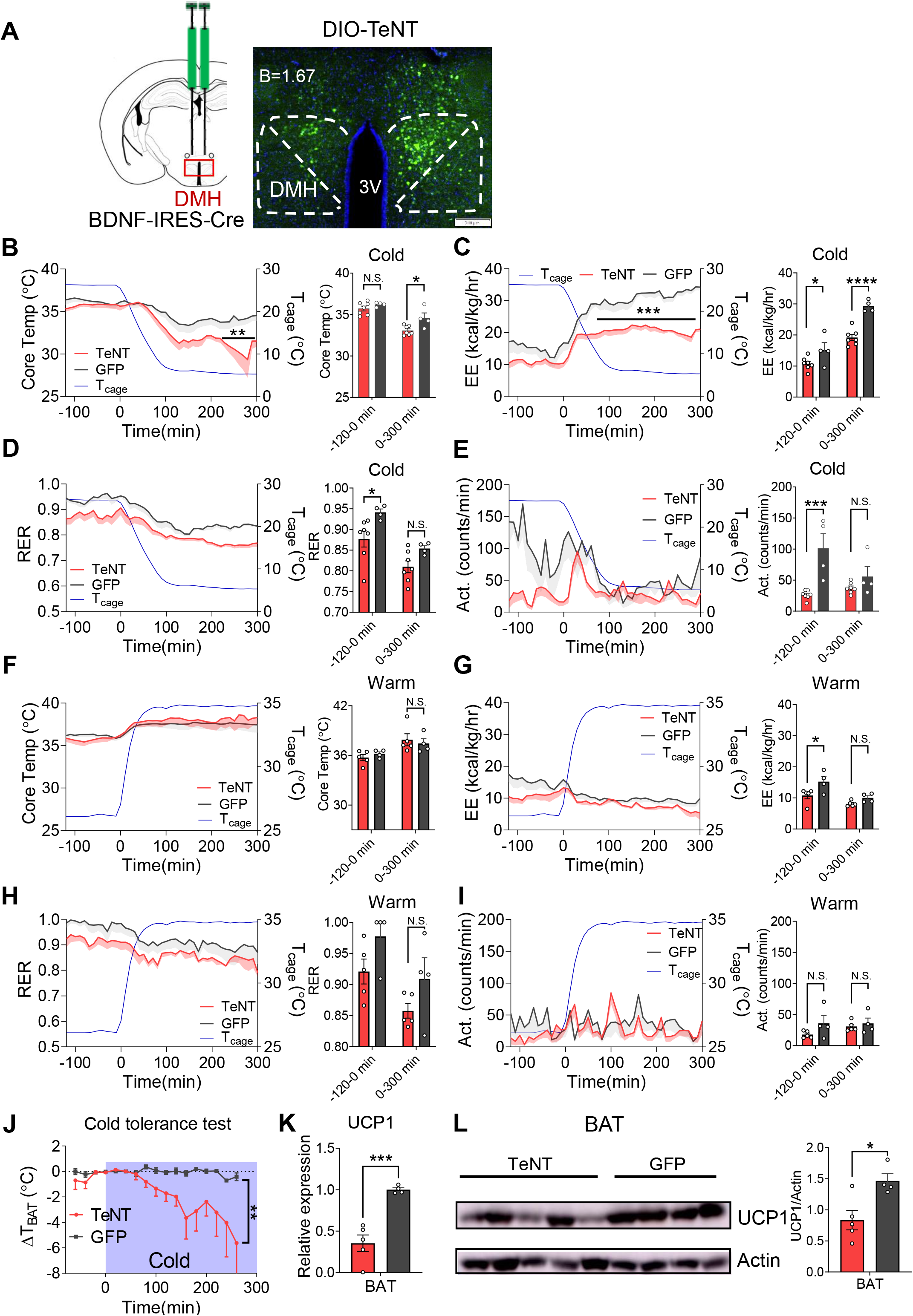
DMH^BDNF^ neurons are required for BAT thermogenesis during cold defense. (**A**) Schematic of virus injection (left) and representative expression of DIO-TeNT-GFP (right) in the DMH of BDNF-IRES-Cre mice. Scale bar, 200 μm. (**B**-**E**) Dynamics of core body temperature (**B**), energy expenditure (**C**), respiratory exchange ratio (RER) (**D**) and physical activity (**E**) in the TeNT (n=7) and control (n=4) mice after cold challenge. For quantification, the bar graph was shown on the right panel. (**F**-**I**) Dynamics of core body temperature (**F**), energy expenditure (**G**), respiratory exchange ratio (RER) (**H**) and physical activity (**I**) in the TeNT(n=5) and control (n=4) mice after warm challenge. (**J**) Change in BAT temperatures during cold tolerance test (CTT; n =6 per group). Blue shadow indicates cold (4°C) duration. BAT temperatures were recorded by a micro-thermosensor implanted in the fat pads (IPTT-300). (**K**) Quantification of BAT *UCP1* mRNA expression level by q-PCR. (**L**) Western blot analysis of UCP1 expression in the BAT. Quantification of the UCP1 band was shown in the right panel. For quantification, the bar graph was shown on the right panel. B, bregma; 3V, 3rd ventricle; DMH, dorsomedial hypothalamus. The *P* values were calculated based on repeated measures two-way ANOVA with Bonferroni’s corrections(**B**-**J**) and two-tailed unpaired t-test (**K**-**L**). Data are presented as mean±SEM; N.S., not significant, *\*P* ≤ 0.05, *\*\*P* ≤ 0.01, *\*\*\*P* ≤ 0.001 and *\*\*\*\*P* ≤ 0.0001.

Based on these defects in cold defense, we hypothesized that BAT thermogenesis would also be affected, since it contributes up to 60% of cold adaptive thermogenesis (***Abreu-Vieira et al. 2015***). We first measured BAT temperature using a small telemetry probe embedded between the two interscapular fat pads. Following cold exposure, BAT temperature was significantly lower in mice lacking DMH^BDNF^ activity compared with controls (***Figure 3J***). Next, we measured changes in the major thermogenin, UCP1. We found significant reductions in both UCP1 mRNA and protein (***Figure 3K-L***), suggesting the BAT thermogenesis was impaired when DMH^BDNF^ neurons were blocked.

### Long-term blocking of DMH^BDNF^ neurons increases body weight and impairs glucose metabolism

We reasoned that impaired BAT thermogenesis in response to long-term blocking of DMH^BDNF^ neurons might impact body weight and glucose metabolism. Indeed, inhibiting these neurons resulted in increased body weight and fat mass compared with controls two months after viral injection (***Figure 4A-B***). This overweight phenotype did not result from increased food intake as the TeNT group and controls consumed the same amount of food one month after viral injection (***Figure 4C***). We continued this line of analysis by measuring glucose clearance and insulin tolerance using the glucose tolerance test (GTT) and insulin tolerance test (ITT), respectively. Intraperitoneal injection of glucose (2 g/kg) quickly increased blood glucose, reaching a peak within 30 minutes (***Figure 4D***). Compared with controls, TeNT mice exhibited higher blood glucose and a larger area under curve, suggesting impairment in the ability to clear blood glucose after neural blocking (***Figure 4D***). Whole-body insulin sensitivity was also significantly lowered (***Figure 4E***). Taken together, these results show that long-term blocking of DMH^BDNF^ neurons reduced BAT thermogenesis resulting in reduced EE and body weight gain.

**Figure 4.**
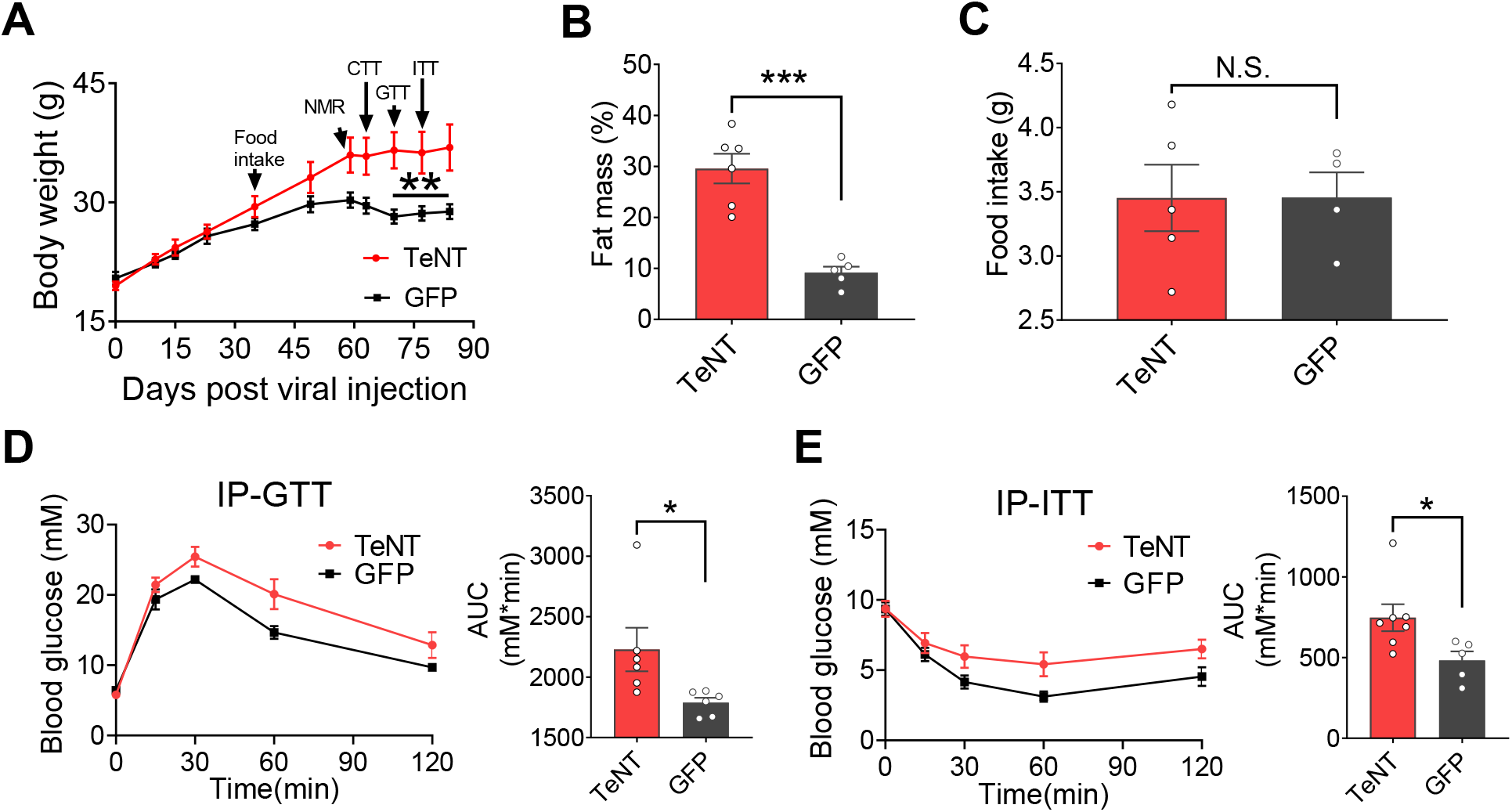
Long-term blocking of DMH^BDNF^ neurons increases body weight and impairs glucose metabolism. (**A**)Body weight of TeNT- and GFP-injected mice after viral injection (n = 7 for TeNT group; n = 5 for GFP group). (**B**) Fat mass of TeNT and control mice 59 days post viral injection measured by NMR. (**C**) Daily food intake of TeNT-and GFP-injected mice 30 days after viral injection. (**D**-**E**) Intraperitoneal tests of tolerance to glucose (IP-GTT) (**D**) and insulin (IP-ITT) (**E**)) in TeNT and GFP mice. For IP-GTT, n=6 per group; for IP-ITT, n = 7 for TeNT group; n = 5 for GFP group. Area under the curve were shown on right panel. The *P* values were calculated based on repeated measures two-way ANOVA with Bonferroni’s corrections (**A**) and two-tailed unpaired t-test (**B**-**E**). Data are presented as mean± SEM; N.S., not significant, *\*P* ≤ 0.05, *\*\*P* ≤ 0.01, *\*\*\*P* ≤ 0.001 and *\*\*\*\*P* ≤ 0.0001.

## Conclusion

These analyses reveal that DMH^BDNF^ neurons function as cooling detectors in the thermoregulatory pathway to control body temperature, EE, BAT thermogenesis, and physical activity, with links to body weight control and glucose metabolism. Given that the human BDNF circuitry has been linked to the development of obesity (***Wen et al. 2012; Gray et al. 2006; Yeo et al. 2004***), our findings provide key mechanistic insights toward understanding the physiology of human obesity.

## Materials and methods

### Experimental animals

All experiments were performed on male adult mice housed under constant temperature (22–25 °C) in a light/dark cycle (light time, 9 PM to 9 AM) with ad libitum food and water. BDNF-IRES-Cre mice were made through the CRISPR/Cas9 genome targeting reported previously (***Luo et al. 2019***). EEF1A1-LSL.EGFPL10 (L10 is a ribosomal resident protein), were gifted by Dr. Jeff Friedman (***Stanley et al. 2013***). All procedures conducted on animals were conformed to institutional guidelines of ShanghaiTech University, Shanghai Biomodel Organism Co., the Animal Facility at the National Facility for Protein Science in Shanghai (NFPS), and governmental regulations.

### Stereotaxic brain injection

We delivered ∼150 nl of AAV virus with a stereotaxic instrument (David Kopf Instruments, #PF-3983; RWD Life Science, #68030) and a pressure micro-injector (Nanoject II, #3-000-205A, Drummond). The fiber optic cables (200 μm in diameter, Inper Inc., China) were tardily implanted and secured with dental cement (C&B Metabond® Quick Adhesive Cement System, Parkell, Japan). The injection coordinate for DMH was calculated according to Paxinos & Franklin mice brain coordinates (3^rd^ edition): -1.27 mm posterior to bregma, 0.27 mm lateral, and 5.20 mm below the skull surface. AAVs (AAV2/9-hSyn-Flex-GCaMP6s-WPRE-SV40pA, AAV2/9-hSyn-DIO-TeNT-GFP, AAV2/9-hSyn-DIO-hM3D (Gq)-EGFP, AAV2/5-EF1a-DIO-EGFP-WPRE-pA) were purchased from Shanghai Taitool Bioscience Co.

### Immunohistochemistry

Mice were anesthetized and perfused transcardially with PBS followed by 50 ml 4% PFA without post-fixation. Brains were dehydrated in sucrose (15% and 30%, two days) at 4°C and sectioned at 40 μm thicknesses on a cryostat microtome (Leica, CM3050s). The brain slices were collected into three sets. For cFos staining, the collected slices were incubated with primary antibodies (rabbit anti-cFos, Synaptic systems, #226003, 1: 10000; Primary Antibody Dilution Buffer, Beyotime#P0262) overnight at 4°C after blocked with the commercialized blocking buffer (Beyotime #P0260). For glutamate and GABA staining, the brain slices were blocked with the blocking solution (10% normal goat serum and 0.7% Triton X-100 in PBS) for 2-h at room temperature and subsequently incubated with primary antibodies (rabbit anti-Glutamate Sigma, #G6642, 1:000 and rabbit anti-GABA, Sigma, #A2052, 1:500) for 6∼8 hours at room temperature. After that, the slices were washed three times in 0.7% PBST and incubated in secondary antibodies (Alexa Fluor 594 conjugated goat anti-rabbit IgG Jackson, #111-585-144, 1: 1000; Alexa Fluor 647 conjugated goat anti-rabbit IgG, Invitrogen, #A21244, 1: 1000,) for 2 hours at room temperature. Then, the brain sections were washed three times in PBST and mounted on coverslips with DAPI Fluoromount-G mounting medium (SouthernBiotech, #0100-20). Images were captured on a Zeiss Z2 Apotome or Leica SP8 confocal microscope or Olympus VS120 Virtual Microscopy Slide Scanning System.

### Calcium fiber photometry

The optical fiber (Inper Inc.) was implanted 100 μm above the viral injection site. For temperature stimuli, the floor temperature was controlled by a Peltier controller (#5R7-001, Oven Industry) with customized Labview code (National Instrument) and was monitored using T-type thermocouples. Floor temperature was converted to a voltage signal and was simultaneously acquired with fluorescence signals in a fiber photometry system (Fscope, Biolinkoptics, China).

Data analysis referred to previous reports (***Zhao et al. 2017; Yang et al. 2020***). To calculate the fluorescence change ratios, we derived the values of fluorescence change (ΔF/F_0_) by calculating (F−F_0_)/F_0_, where F_0_ is the baseline fluorescence signal averaged in a time window before events (temperature changes) indicated in the figure legends. We used GraphPad Prism 8 (GraphPad) to plot the fluorescence changes (ΔF/F_0_) with different stimuli.

### Bodyweight and body composition analyses

Bodyweights were continuously measured for all animals before mice sacrificing. Body composition was determined using a Minispec whole-body composition analyzer (Burker Minispec CMR LF50).

### Metabolic measurement

The energy expenditure, respiratory exchange ratio, body temperature, and locomotor activity were monitored by the Comprehensive Lab Animal Monitoring System with Temperature Telemetry Transmitter (CLAMS; G2 E-Mitter). Temperature transponders were implanted into the abdominal cavity 3–5 days before testing. Mice were adapted in the metabolic chambers for two days before giving temperature stimuli or CNO (i.p., 2.5 mg/kg body weight, ENZO, #BML-NS105-0025).

### Cold tolerance test (CTT)

TeNT (as to their controls) injected mice were exposed to a 4-h cold exposure and recording interscapular brown adipose tissue temperature simultaneously to assess BAT thermogenesis capacity, which is defined by cold tolerance test (CTT) in our study. The IBAT temperature was measured three times for average using subcutaneous implanted thermal probes (IPTT-300, Biomedic Data Systems). The probes were implanted at the midline in the interscapular region at least one week before testing.

### Glucose and insulin tolerance test (IP-GTT / IP-ITT)

For IP-GTT, fasting blood glucose was measured after 16-h fasting. Then mice were intraperitoneally injected glucose (2 g/kg). For IP-ITT, mice were fasted for 4 hours and challenged with an intraperitoneal injection of insulin (0.75 IU/kg). Blood glucose was measured at 0, 15, 30, 60, 90, and 120 min using a hand-held glucometer (Accu-Chek Performa Connect, Roche, Switzerland).

### Real-time PCR

Total RNA was isolated from iBAT using trizol (Sangon, #B511311-0100). Total RNA (1 µg) was reverse-transcribed with PrimerScript^™^ RT Master Mix (Takara, #RR047A) after DNase I treatment (Takara, #RR047A). PCR was performed in triplicate using an SYBR Green PCR kit (Takara, #RR820A) in a 10 µl volume. *UCP1* expression levels was calculated based on the 2^−ΔΔCT^ method. *Actb* was used as a reference gene.

### Western blot analysis

IBAT was lysed in RIPA buffer for protein extraction. Lysates were separated by 10% SDS-PAGE and transferred to a Nitrocellulose membrane (Millipore, #HATF00010). The membranes were incubated with anti-UCP1 (rabbit polyclonal, 1:1000; BBI #D262447-0025) or β-Actin (rabbit polyclonal, 1:5000; Beyotime #AF5001) at 4°C for 12 h and washed three times in 0.1% TBST, then sequential incubating with secondary antibodies (Goat anti-rabbit polyclonal IgG conjugated with HRP, 1:5000; BBI; #D110058-0100) at room temperature for 2 hours. Protein levels were quantified using ImageJ software.

### Statistical analyses

Excel and GraphPad Prism 8 were used to plot data and calculate statistical significance. All data are shown as mean ± SEM. *P*<0.05 was significant.

## Acknowledgments

We thank Dr. Xiaoming Li and the Molecular Imaging Core Facility (MICF) of School of Life Science and Technology, ShanghaiTech University for Microscopic imaging; Dr. Ying Xiong and the Molecular Cellular Core for slices and staining. This study was funded by National Nature Science Foundation of China (H. Sun, 31900707; W. Shen, 31771169 and 91857104), China Postdoctoral Science Foundation (H. Sun, 2019M651616), Shenzhen-Hong Kong Institute of Brain Science-Shenzhen Fundamental Research Institutions (W. Shen, NYKFKT20190017), and Shanghai Science and Technology Committee of Shanghai City (19140903800). We thank the staff members of the Animal Facility at the National Facility for Protein Science in Shanghai (NFPS), Zhangjiang Lab, China, for providing technical support and assistance.

